# Species traits predict extinction risk across the Tree of Life

**DOI:** 10.1101/2020.07.01.183053

**Authors:** Filipe Chichorro, Fernando Urbano, Dinarte Teixeira, Henry Väre, Tiago Pinto, Neil Brummitt, Xiaolan He, Axel Hochkirch, Jaakko Hyvönen, Lauri Kaila, Aino Juslén, Pedro Cardoso

## Abstract

Biodiversity is eroding at unprecedented rates due to human activity^1^. Species’ trajectories towards extinction are shaped by multiple factors, including life-history traits^2^ as well as human pressures^3^. Previous studies linking these factors to extinction risk have been narrow in their taxonomic and geographic scope^4^, thus limiting the ability for identifying global predictors. We studied the relation between 12 traits and the extinction risk of almost 900 species representing 15 groups across the tree of life (vertebrates, invertebrates and plants) at a global scale. We show that threatened species share narrow habitat breadth, poor dispersal ability, low fecundity, small altitudinal range, and are affected by a large human footprint. Other traits either show contrasting responses among groups (body size, offspring size, and change in human footprint), or relations were found for only a limited number of taxa (generation length, diet breadth, microhabitat). Our study suggests that in the absence of data on the precise distribution and population trends of species, traits can be used as predictors of extinction risk and thus help guide future research, monitoring and conservation efforts.

## Main

We are currently facing the sixth mass extinction at the planetary scale. Species are becoming extinct at rates 1000 to 10000 faster than background extinction rates^1^. Not only species, but functions they provide and that benefit humanity are at risk, with unpredictable consequences towards our own well-being. And yet, we are mostly unaware of what species are most at risk and why, with many becoming extinct even before description: the Centinelan extinctions. This lack of knowledge can be partly circumvented if we know which characteristics, or traits, are common to endangered species and which allow species to be resilient to anthropogenic change.

The vulnerability of species to extinction largely depends on their life-history strategies (intrinsic traits), and biotic and abiotic conditions species face (extrinsic traits)^2,3,5^. All studies linking the extinction risk of species to intrinsic and extrinsic factors have focused, however, on few species or narrow geographic ranges. Due to societal and knowledge biases^6,7^, well-studied groups include vertebrates, namely mammals^2,3^ and birds^8,9^ and the best-known region is the Palearctic realm^4,10^. The relations between traits and extinction risk across the tree of life have never been analyzed at a global taxonomic and geographic scale.

### A global trait analysis

Here, we compiled a dataset of 12 traits commonly related to extinction risk (Table S1): body size, offspring size, fecundity, generation length, diet breadth, trophic level, dispersal ability, microhabitat, habitat breadth, altitudinal range, human footprint within the species range as of 2009^11^, and the relative change in human footprint over a period of 16 years (1993-2009)^11^. Traits were quantified for 874 species in five groups of each of vertebrates (mammals, birds, reptiles, amphibians, and fishes), invertebrates (dragonflies, butterflies, grasshoppers, spiders and snails) and plants (bryophytes, ferns, gymnosperms, monocots and legumes) (Table S2-4). Each of the 15 taxonomic groups included 10 species in each of six biogeographic realms, five threatened and five non-threatened, as long as data on extinction risk was available, namely global assessments in the International Union for the Conservation of Nature (IUCN) Red List of Threatened Species (see methods, Table S5). We used these data to identify global predictors of extinction risk across taxa and space.

For all groups, we first standardized trait values to ensure comparability. We then inspected the existence of relationships between traits with pair plots and Spearman rank correlations (Fig. S1). As no strong correlations were found we used all traits in subsequent analyses. We modelled the extinction risk as a binary response variable (threatened versus non-threatened following the IUCN Red List categories: threatened: EX, CR, EN, VU and NT, non-threatened: LC; note that our grouping is different from the usual for IUCN). Significant differences between threatened and non-threatened species were tested both within and for all taxonomic groups. Within groups, significant differences were detected with null models, by comparing the mean and the standard deviation of trait values of threatened species with a distribution of simulated data, sampled across all the possible values for that trait and group. We applied Bayesian mixed models to detect significant relationships between traits and extinction risk across taxa. The mixed models were used to relate threat status against each trait, while controlling for the non-independent effects of taxonomy (using the taxon grouping as a random effect in the models). We inferred significance in either positive or negative relationships between the extinction risk status and each trait when 95% of the posterior distributions of the estimates were not intercepting the zero value. We also related the geographical range size of species to extinction risk (Fig. S2), but we excluded it from further analyses because this trait is used to determine extinction risk in most IUCN Red List assessments. Moreover, range size itself may often not be the driver, but a consequence of trajectories towards extinction, such as population size and trend. Range and population size could only be used without circular reasoning if pre-disturbance values were known, which is almost invariably not the case.

### Predicting extinction risk

Five traits were found to be consistently (similar sign across groups) and significantly (p < 0.05, or almost significantly, p < 0.1) related with extinction risk (Table 1): habitat breadth, dispersal ability, fecundity, altitudinal range, and human footprint.

**Table 1:** Results of the MCMCglmms relating the value of each trait with extinction risk.

| Trait | Sign | pMCMC | N species |
| --- | --- | --- | --- |
| Body size | 0 | 0.666 | 825 |
| Offspring size | 0 | 0.182 | 438 |
| Fecundity | - | 0.0467 | 333 |
| Generation length | 0 | 0.384 | 340 |
| Diet breadth | 0 | 0.77689 | 214 |
| Trophic level | 0 | 0.689 | 266 |
| Dispersal ability | - | 0.00543 | 335 |
| Microhabitat | 0 | 0.802 | 545 |
| Habitat breadth | - | <0.0001 | 874 |
| Altitudinal range | - | 0.0653 | 314 |
| Human footprint | + | 0.000857 | 561 |
| Change in Human footprint | 0 | 0.73131 | 561 |

The association between habitat breadth and extinction risk was negative, highly significant (Table 1) and found across all taxa (Fig. 1). Species occurring in a narrower range of habitats have fewer opportunities to expand to and survive in alternative suitable living conditions and are consequently more likely to be threatened^12,13^. In fact, habitat breadth, together with geographical range size and abundance, is one of the three classical dimensions of rarity^14^. In a previous meta-analysis^4^ habitat breadth was the only factor, besides geographical range size, that was consistently found to correlate with extinction risk. This trait should be very relevant in the face of generalized natural ecosystem destruction with consequent habitat loss for numerous species. With increasing levels of habitat loss occurring across all biomes, species that adapt to alternative habitat types will inevitably fare better.

**Figure 1:**
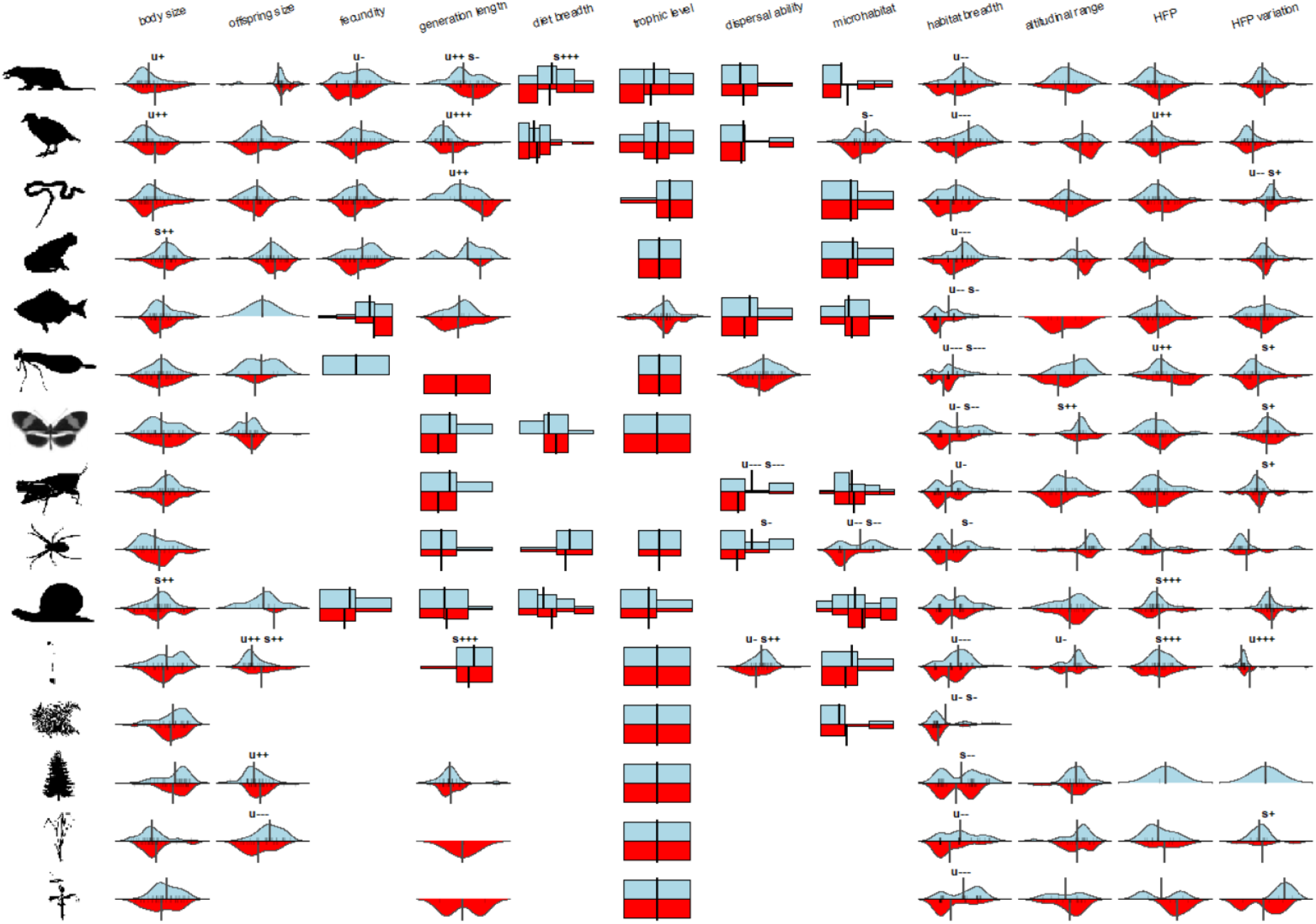
Beanplots of density of trait values between non-threatened (blue, upper side) and threatened (red, lower side) species. Small vertical bars represent one species’ value; darker bars indicate several species with the same trait value. The large vertical bar is the mean trait value. Null models show whether the mean (u) or standard deviation (s) of trait value of threatened species is higher (+++, ++, +) or lower (−−−, −−, −) than expected. Significance codes: +++ or −−− x < 0.01; ++ or −− 0.01 <= x < 0.05; + or - 0.05 <= x < 0.1.

We identified a negative association between dispersal ability and extinction risk (Table 1), indicating that species with poor dispersal ability are more likely to be at risk than those dispersing easily. The pattern was common across tested taxa (mammals, birds, dragonflies, grasshoppers, spiders, and bryophytes) but stronger in grasshoppers, spiders and bryophytes (Fig. 1). In the face of localized threats, species with a better capacity to colonize new areas have a lower risk of extinction^15,16^. Consequently, species groups with high dispersal capability, such as birds and dragonflies, often have a lower number of threatened species compared to other groups. In addition, shifts in species’ distributions caused by climate change are likely to exacerbate the extinction of poor dispersers^17,18^. This trait might be very relevant in predicting which species will be able to adapt to increasing levels of fragmentation of natural habitats. As fragmentation is one of the consequences of habitat loss, its effects are increasing at global scales, and having good dispersal ability might prove crucial to the survival of many species.

The model relating fecundity to extinction risk included mostly data from vertebrates (mammals, birds, reptiles, amphibians, and fishes) and one invertebrate group (snails, table S3). Fecundity was negatively associated with extinction risk (Table 1), indicating that species with lower reproductive output are more at risk, with the strongest signal for mammals (Fig. 1). Populations of species with low reproductive output are presumed to decline in the face of demographic threats, due to diminished capacity to compensate for higher mortality rates^19^. Mammal families with higher proportions of species threatened by hunting and fishing have smaller litter sizes^20^. Although our model for fecundity includes no plant species due to lack of data, it is likely that species with lower reproductive output are also more threatened in plants^21^. Species with higher reproductive output will probably fare better in the future independently of any particular threat.

Altitudinal range, often a measure of climatic tolerance, was marginally negatively correlated with extinction risk (Table 1). The modest significance of this trait could be a consequence of the small number of species for which data were available. The signal was stronger for bryophytes and, to some extent, butterflies (Fig. 1). Species with lower climate tolerance have fewer chances to be able to exploit new ranges for survival, and thus confronting higher extinction risk^22^. This trait in particular might be very relevant during the current climate emergency, as it might indicate which species will be able to adapt in the future to changing temperature and precipitation patterns.

The human footprint index was positively related to extinction risk (Table 1), indicating that species with higher mean human pressure within their ranges are more likely to be threatened. The pattern was consistent across taxa, but stronger in birds, dragonflies, snails and bryophytes (Fig. 1). This result was expected and reflects the fact that most organisms, independently of their traits, are sensitive to human pressure. Synanthropic or bred/cultivated species are obvious exceptions, benefiting from human pressure across their range^23^.

While the previous five traits were found to be global predictors of extinction risk, some were found to differ in their signal according to the species group (Supplementary discussion). They either showed contrasting responses between groups (body size, offspring size, and change in human footprint) or relations were found for only a limited number of taxa (generation length, diet breadth, microhabitat). In some cases a similar trait might in fact reflect different competitive advantages depending on the group, such as larger body sizes being targeted by hunting in the case of mammals and birds, but constituting a competitive advantage for many plant taxa^4^. In other cases, either data are missing or have little contrast for some taxa, preventing general trends to be found. Future studies with more data might help clarify and further support some of the trends already found.

### Future directions

In this work, we were able to study the relationships between all the main traits found in the past to influence extinction risk, and the threat level of species covering numerous branches of the tree of life from many parts of the world. Notable exceptions are fungi and marine taxa, for which knowledge is scarcer. Inevitably, there are still gaps in both the knowledge available on traits (e.g. fecundity, dispersal ability) in some taxa and of geographic coverage (mainly tropical species) in others. Yet, our results are not only robust, but also generalizable for a wide spectrum of terrestrial organisms.

Our study supports the view that extinctions do not affect species randomly, but extinctions are rather mediated by species traits^23^. We show that across the tree of life, species with a higher extinction risk are those with narrow habitat and climatic niches, poor dispersal capacity, and low fecundity. On top of this, the presence of human activity increases the probability that the species become threatened. These results emphasize two different aspects of extinction: firstly, with species extinctions, we might not only lose species but also their particular functions in ecosystems, which in turn, could lead to further co-extinctions. Secondly, high human impact on ecosystems is generally not compatible with species survival.

With the understanding of key biological factors contributing to species vulnerability, we will be able to identify species that are more prone to extinction, even in the absence of data that are most commonly used but often unavailable, such as geographic range size or current population trends. These two factors have been considered as the most important for extinction assessments according to the IUCN Red List criteria. Yet, often they are unknown or biased, with figures above 50% Data Deficient species reported for invertebrates, which represent the vast majority of species^24^. Using traits as surrogates for extinction risk will help reduce this knowledge gap, allowing prioritization of future research, monitoring and conservation efforts.

## Supplementary materials

### Methods

Because our goal was to find general trends, we selected 1) species belonging to a variety of taxonomic groups covering most of the multicellular tree of life (vertebrates, invertebrates and plants), and 2) traits that were generalizable across taxa, not considering others that would be specific for some groups (e.g., brain size).

#### Selection of species

As a first step in selecting the species, we identified a range of taxa that would capture a wide range of life-histories and geographical locations. We restricted our species pool to the species already assessed for the global IUCN Red List of Threatened Species™ (www.iucnredlist.org), excluding those that were assessed as Data Deficient. We also restricted our analysis to the multicellular branch of the tree of life. Since very few assessments of fungi exist, we also excluded these. Finally, we restricted the analysis to terrestrial and freshwater species given that the few marine species assessed would require different stratified sampling and analyses. We therefore chose five vertebrate, five invertebrate, and five plant groups. Vertebrate groups comprised “Mammals” (Class: Mammalia), “Birds” (Class: Aves), “Reptiles” (Class: Reptilia), “Amphibians” (Class: Amphibia) and freshwater “Fishes” (Class: Actinopterygii). The invertebrate groups comprised “Dragonflies” (Order: Odonata, including damselflies), “Butterflies” (Suborder: Rhopalocera), “Grasshoppers” (Order: Orthoptera), “Spiders” (Order: Araneae), and land “Snails” (Class: Gastropoda). In the selection of the plants we followed the recent baseline study^25^ with the following embryophytes (land plants): “bryophytes”, excluding hornworts (Divisions: Bryophyta and Marchantiophyta), “pteridophytes” (Classes: Lycopodiopsida, Polypodiopsida), “Gymnosperms” (Classes: Pinopsida, Cycadopsida, Gnetopsida), “Monocots” (Class: Liliopsida), and finally the “Legumes” (Order: Fabales) serving as a representative of the largest group of plants, eudicots (Table S5).

To guarantee global spatial representativeness of the dataset, we selected, whenever possible, 10 species per group from each of six biogeographic realms (Table S5). Of the 10 species, we randomly selected from the global IUCN database equal numbers of threatened (one of each of Near Threatened, Vulnerable, Endangered, Critically Endangered or Extinct) and non-threatened (five Least Concern) species.

As not all taxa have global coverage in the IUCN Red List, we had to restrict our dataset to smaller regions in the case of butterflies, grasshoppers, spiders, snails, and bryophytes. All species of spiders, snails and bryophytes were selected from Europe, due to very low numbers of assessments from other geographical realms. For grasshoppers, most assessments came from the Afrotropics and Palearctic, and therefore 30 species were selected from each. In butterflies, no Nearctic species were included due to unavailability of data from that group by the time we made the selection (IUCN version 2018-2), and very few from the Indo-Malay region were included. In total, our dataset included 874 species.

#### Selection of traits

As predictors, we selected intrinsic and extrinsic traits of species whose relationship with extinction risk has been hypothesized and tested in previous studies for some taxonomic groups (Table S1) but excluded traits that are specific to a few taxa only (e.g. brain size). Intrinsic trait data were compiled from the literature, including existing trait databases, and in some cases also measurements of photographs of pinned specimens (usually the holotype or paratype of species) available online.

Different taxonomic groups differ in their life-history and ecological strategies. Therefore, for each intrinsic trait, we selected trait “proxies” (Table S3), which are analogous traits^26^ with the same function across taxa but measured differently. The choice of proxies depended on the suitability of the trait as a proxy (e.g., body length is a better proxy of body size than body mass in birds, due to large variation within a species between seasons), and on the availability of data for that trait (e.g. dispersal ability of birds and mammals being a binary trait distinguishing migratory and/or nomadic species from those not, an ordinal trait reflecting the propensity to balloon in spiders and a continuous trait of seed size in plants).

To measure the human footprint pressure and the change in human footprint within each species’ range, we used recently constructed 1km^2^ resolution raster maps of human footprint available for the years 1993 and 2009^11^. In these maps, each raster cell is characterized by a score of cumulative human footprint pressure, ranging from 0 (no human impact) to 50 (very high human impact). The score of a grid cell is a function of the presence and/or magnitude of eight types of pressures: the extent of built environments, human population density, electric infrastructure, crop lands, pasture lands, roads, railways and navigable waterways^11^. To estimate the mean human footprint of 2009 across a species’ range, we averaged all grid cell values within each species polygon maps, retrieved from IUCN (see below). To estimate the change in human footprint, we first constructed a map of the differences between 2009 and 1993, with positive values indicating a positive change in human footprint (more human impact in 2009 compared to 1993) and negative values indicating negative change, and then averaged the scores across species’ ranges. The species’ range maps were obtained from the IUCN Red List of Threatened Species. We only included maps with the following origin, presence, and seasonal descriptors: “Native” or “Reintroduced”; “Extant”, “Probably extant”, or “Possibly extant”; and “Resident”, “Breeding season”, “Non-breeding season” or “Seasonal presence uncertain”.

When no trait data were available for the species, we used either the value of a closely related species or the genus or family average; this latter approach was used when values for other taxa were available in online trait databases. Genus and family averaging were never performed for binary data, habitat breadth, altitudinal range, human footprint, change in human footprint, and geographical range size variables.

Some groups lacked data completely for some traits, such as fecundity and offspring size for dragonflies and spiders, and diet breadth for reptiles and amphibians. Trophic level was known for all species but in some groups the trophic position resolution was finer (fishes), while for some others it was coarse or invariant (dragonflies, spiders, plants).

Because offspring size is highly correlated with body size, we used instead a relative metric of offspring size: the residuals of a regression between log(offspring size) and log(body size) within all groups. Because the altitudinal range is often related to the geographical range size of species, we used the residuals of a regression between log(altitudinal range) and the log(geographical range size).

The compiled dataset included data for 94% and 99% of the species for body size and habitat breadth, respectively (Table S2). Particularly for invertebrate and plant species, data availability for some traits was low, including offspring size (mean 50%, range 0% to 100%), fecundity (38%, 0-100%), generation length (39%, 0-100%), diet breadth (24%, 0-100%), or dispersal ability (38%, 0-100%). The human footprint and the change in human footprint were available for 64% of species, since species’ maps are available for many of the species on the IUCN website.

#### Trait transformation and standardization

We log-transformed count data (e.g. number of habitat types, number of diet types eaten), and continuous data (body length, number of offspring), except the dispersal ability of dragonflies and bryophytes, and residuals of offspring size and altitudinal range, since these traits were already log-transformed when estimating their values. This ensured that the distribution of trait values followed a near-normal distribution without observations spread far away from the main density of trait values. For extinct species, which have geographical range sizes of 0 km^2^, we replaced these 0’s with 0.1, so that log-transformation of these data points was possible. Likewise, altitudinal range values lower than 10m were converted to 10m.

Within a given trait, the units and measurement scales were different across groups and it was necessary to transform these data to guarantee comparability between taxa. All trait values within groups were subject to a z-transformation, which includes rescaling of data (by dividing each data point by the standard deviation) and recentering by subtracting the mean value from each observation. This type of scaling preserves the mean and standard deviation of each trait.

#### Statistical analysis

We first modelled each trait separately (univariate models) per group. To check for significant relationships between each trait distribution and extinction risk within groups, we ran null models. The null model compared the mean and the standard deviation of the trait values of the threatened species with the mean and standard deviation of 1000 null expectations when extracting the same number as threatened species from the complete pool of threatened plus non-threatened. A deviation from the null expectation was considered to have been met when either the mean or the standard deviation of the threatened species were lower or higher than the 2.5^th^ or the 97.5^th^ percentiles of the null distribution, in which case a significant negative or significant positive deviation was annotated respectively.

To test the relation between individual traits and extinction risk across taxa we used generalized linear mixed effect models (GLMMs), in which we controlled the effect of taxonomy by allowing a random intercept and random slope dependent on the taxonomic group. Since our response variable was binary, our GLMM consisted of a logistic regression. The GLMMs were modelled within a Bayesian framework, using Monte Carlo Markov Chains with R package MCMCglmm^27^. We used the default priors of package MCMCglmm, which are weak priors. We ran simulations with 50,000,000 iterations, excluding the first 1,000,000 (burn-in). To ensure good mixing of chains, we only saved every 1000^th^ iteration (thinning). With these parameters, we observed good mixing of chains and thus, good convergence of posterior parameters’ distributions.

The random terms in the model add a new assumption to the overall model, which is that taxonomic groups are sampled from a larger population of possible taxonomic groups and that the intercepts and slopes of each group follow a normal distribution of intercepts and slopes around the population means of the intercept and slope. The association between the trait and the probability of being threatened was considered to be strong when 95% of the posterior distribution of a trait was not intercepting zero, and moderate for 90%. All statistical analyses were performed in R version 3.6.1^28^. Beanplots were done with package beanplot^29^ and pairwise plots with packages ggplot2^30^ and GGally^31^.

## Supplementary discussion

The distribution of body size values differed between threatened and non-threatened species in birds, amphibians, snails, and marginally in mammals (Fig. 1). Overexploitation might explain the trends for mammals and birds, as larger species are direct targets of hunting^20,32^. Particularly for mammals, as size increases the importance of life-history traits in determining extinction risk increases in relation to extrinsic traits^33^. Larger species within these generally large body-sized taxa (compared with, e.g., insects) also require more resources and these might quickly dwindle to unsustainable levels^33^. In amphibians and snails, the standard deviations of body size values of threatened species were significantly greater than the null expectation (Fig. 1), indicating that a larger proportion of threatened species occur at both ends of the body size distribution for these groups. The conservation status of small body-sized organisms could be explained by a particular set of life-history traits that may predispose them to naturally restricted range size and narrow habitat breadth^34^.

After accounting for body size, the relative offspring size of organisms showed mixed signals across taxa, with the only significant values being found for plant groups, albeit with opposing signals (Fig. 1). In bryophytes and gymnosperms offspring size was positively correlated with extinction risk, while for monocots this relationship was negative. The size of an offspring in relation to the body size is an indication of the investment in reproduction. The trade-off is between small and numerous, or large and scarce. Larger species tend to invest more in large offspring, to compensate for higher mortality during a very long juvenile stage^35^. However, when environmental conditions change rapidly, investing in only a few descendants might be a bad option due to low variability under unpredictability. For monocots the negative relation might be because 19 species in the dataset are orchids, which are characterized by the smallest seeds among plants^35^. Orchids have very specific requirements for survival (their seeds require the presence of mycorrhizal fungi to germinate) and therefore this relationship might be spurious and phylogenetically driven, even if orchids are in fact generally in higher threat categories than most other plant groups.

The change in human footprint indicates the rate of increase or decrease of human footprint in species’ ranges, with higher positive values showing a greater increase in impact, and higher negative values showing a greater decrease in impact. Its influence on extinction risk was diverse, with no congruent pattern across taxa (Fig. 1). Threatened bryophytes were characterized by significantly larger positive changes in human footprint values than non-threatened bryophytes, while threatened reptiles were characterized by significantly larger negative changes than non-threatened reptiles. This might be due to recent impacts leading to large differences in the index that are still to be reflected in species populations. The absence of a global trend indicates that the magnitude, rather than the rate of change in human footprint is related to extinction risk.

Generation length had a very clear relationship with extinction risk for mammals, birds and reptiles (Fig. 1). Just as with fecundity, species with delayed life-cycles are more likely to be more threatened. For organisms with slower life-cycles it takes longer to recover from low population numbers in the face of demographic troughs^19^. The weak or nonexistent effect seen in invertebrates or plants taxa may be due to lack of contrasts in data (either few data available, available just as ordinal values or showing low natural variability).

The mean diet breadth between threatened and non-threatened species did not vary across groups (Fig. 1). However, the range of values differed between threatened and non-threatened mammals, indicating that threatened species occurred at both ends of the range in diet breadth. As for seed size, phylogeny might be playing a role, as diet specialists have been shown to be more at risk within bats^36^, but not within artiodactyls^37^ for example. Further studies including the phylogenetic relations of species would help clarify any general effects of diet breadth on extinction risk. As comprehensive phylogenetic trees are currently available for only some of the taxa we studied this is not possible as of yet.

Microhabitat was a significant factor only for spiders (Fig. 1); spiders occurring at higher vertical strata are less threatened. This effect seems to be due to the presence of organisms with higher capacity for ballooning in this stratum. As ballooning depends on, first, building the right kind of silk strands (more commonly found in web weavers) and second, finding the right place to take off (usually at higher heights) spiders living on trees and other vegetation are often more prone to balloon than those living at ground level^38^. Microhabitat therefore determines to a certain point dispersal ability and consequently extinction risk in spiders.

## Supplementary tables

**Table S1:**
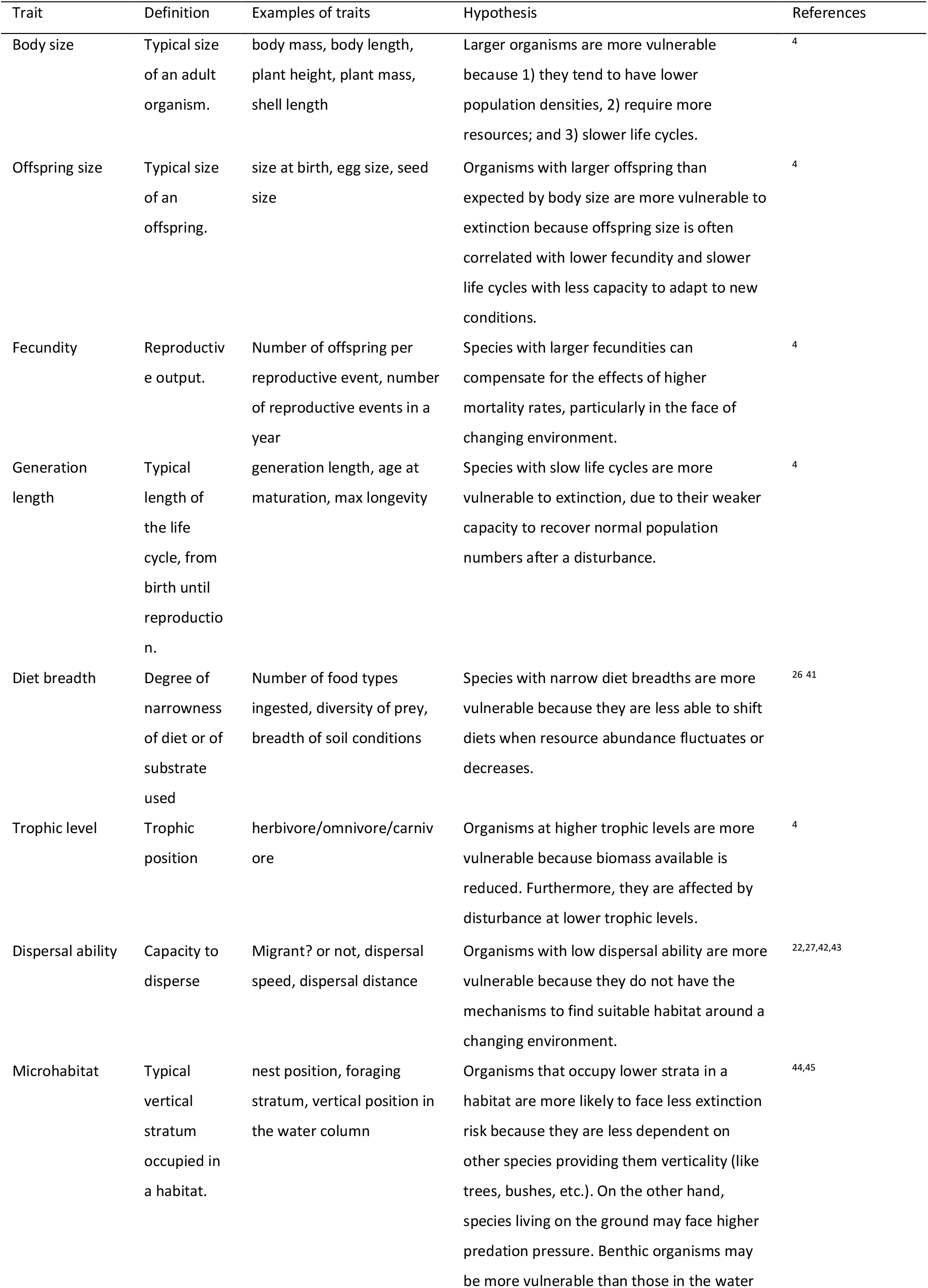
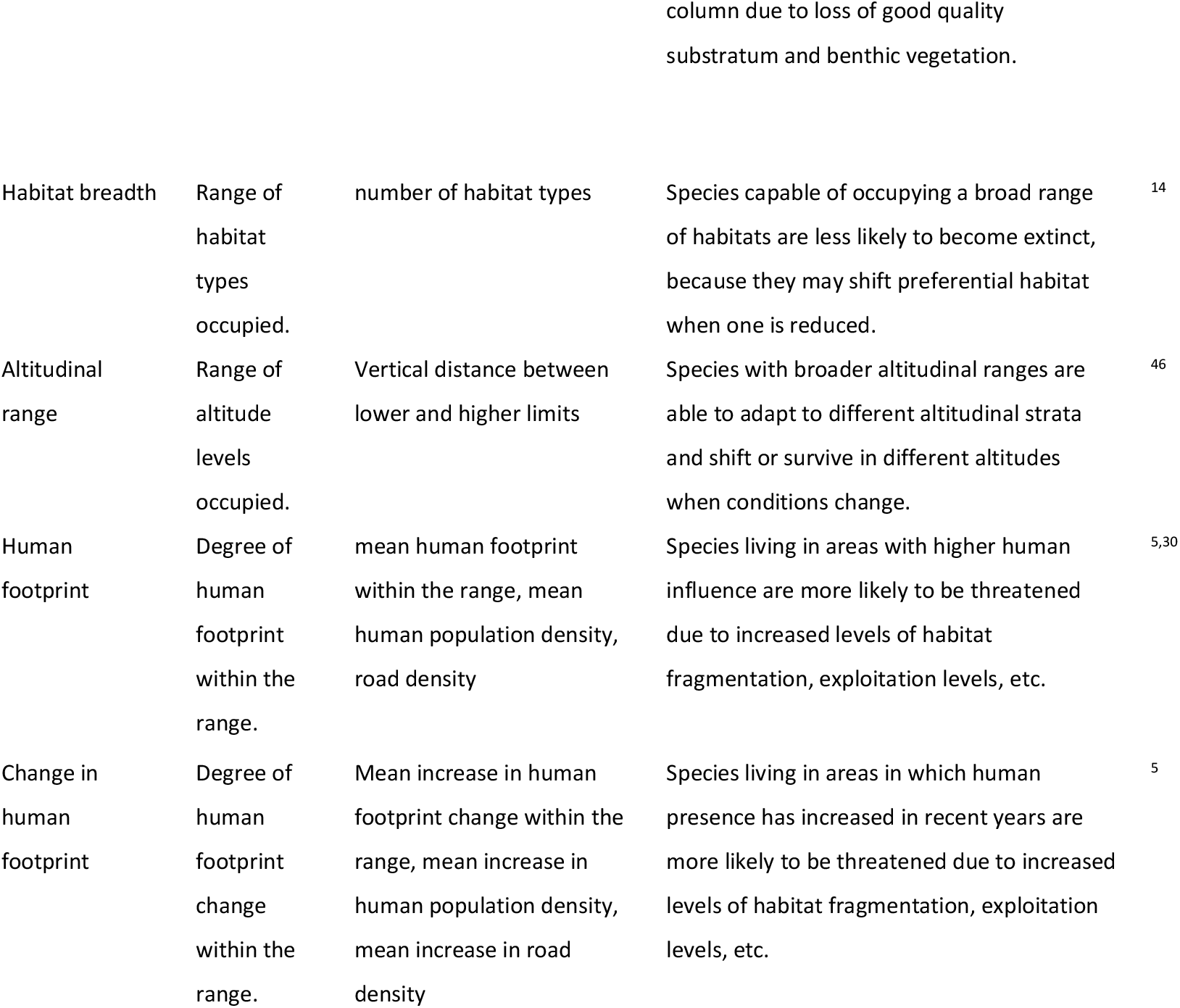
Traits studied, definition, examples, and hypotheses.

**Table S2:** Number of species for each trait in each taxonomic group.

|  | Body<br>size | Offspring<br>size | Fecundit<br>y | Generation<br>length | Diet<br>breadth | Trophic<br>level | Dispersal<br>ability | Microhabitat | Habitat<br>breadth | Altitudinal<br>range | Human<br>footprint |
| --- | --- | --- | --- | --- | --- | --- | --- | --- | --- | --- | --- |
| *Total* | 842 | 415 | 334 | 344 | 214 | 743 | 335 | 551 | 874 | 314 | 562 |
| Mammals | 60 | 59 | 58 | 58 | 59 | 60 | 35 | 60 | 59 | 16 | 55 |
| Birds | 59 | 0 | 58 | 59 | 60 | 60 | 60 | 60 | 60 | 17 | 50 |
| Reptiles | 60 | 43 | 60 | 11 | 0 | 45 | 0 | 44 | 59 | 15 | 51 |
| Amphibians | 60 | 60 | 60 | 6 | 0 | 18 | 0 | 57 | 60 | 21 | 55 |
| Fishes | 60 | 2 | 56 | 9 | 0 | 59 | 29 | 60 | 60 | 5 | 51 |
| Dragonflies | 60 | 5 | 1 | 1 | 0 | 60 | 60 | 0 | 60 | 7 | 34 |
| Butterflies | 59 | 47 | 0 | 10 | 17 | 60 | 0 | 0 | 60 | 19 | 44 |
| Grasshoppers | 51 | 0 | 0 | 5 | 0 | 0 | 60 | 60 | 60 | 27 | 56 |
| Spiders | 39 | 0 | 0 | 39 | 39 | 39 | 39 | 39 | 39 | 36 | 33 |
| Snails | 60 | 9 | 41 | 39 | 39 | 42 | 0 | 59 | 60 | 29 | 56 |
| Bryophytes | 52 | 46 | 0 | 60 | 0 | 60 | 52 | 59 | 60 | 36 | 57 |
| Ferns | 50 | 0 | 0 | 0 | 0 | 60 | 0 | 53 | 60 | 0 | 0 |
| Gymnosperms | 58 | 57 | 0 | 44 | 0 | 60 | 0 | 0 | 60 | 32 | 1 |
| Monocots | 60 | 41 | 0 | 1 | 0 | 60 | 0 | 0 | 59 | 32 | 14 |
| Legumes | 54 | 46 | 0 | 2 | 0 | 60 | 0 | 0 | 58 | 22 | 5 |

**Table S3:** Table of proxies used for each trait. A “−” indicates traits for which we could not find enough data/proxies.

| Group | Body size | Offspring size | Fecundity | Generation length | Diet breadth | Trophic level | Dispersal ability | Microhabitat |
| --- | --- | --- | --- | --- | --- | --- | --- | --- |
| Mammals | Adult body mass (g) | Neonatal mass (g) | Litter size (no. of offspring) | Maximum longevity (years) | Number of food types | 1 = herbivore; 2 = omnivore; 3 = carnivore | 1 = not a migrant; not nomadic; 2 = migrant or nomadic | Foraging stratum: 1 = marine; 2 = ground level, including aquatic foraging; 3 = scansorial; 4 = arboreal; 5 = aerial; categories from Elton traits database<sup>47</sup> |
| Birds | Body length (cm) | Egg volume (mm<sup>3</sup>), estimated from egg height and diameter using the Hoyt equation<sup>48</sup> | Clutch size (no. of offspring) | Generation length (years) | Number of food types | 1 = herbivore; 2 = omnivore; 3 = carnivore | 1 = Not a migrant; 2 = altitudinal/full migrant | Index of foraging verticality from 0 (prevalence of foraging below water) to 1 (prevalence of foraging well above vegetation or other structures). Adapted from Elton traits database<sup>47</sup> |
| Reptiles | Adult body mass (g) | hatchling snout-vent length (mm) | Clutch size (no. of offspring) | Generation length (years) | - | 1 = herbivore; 2 = omnivore; 3 = carnivore | - | Verticality: 1 = ground level (ground dwelling, among rocks, freshwater, leaf litter); 2 = upper level |
| Amphibians | Snout-vent length (mm) | Offspring size (mm) | Clutch size (no. of offspring) | Age at sexual maturity (years) | - | 1 = carnivore | - | Verticality: 1 = exclusively ground level/aquatic; 2 = arboreal or arboreal and/or aquatic and/or terrestrial) |
| Fishes | Body length (cm) | egg diameter (mm) | Minimum population doubling time: 1= more than 14 years; 2= 4.5-14 years; 3 = 1.4-4.4 years; 4 = less than 15 months | Generation length (years) | - | trophic position (average $\delta^{15}\text{N}$ signature) | Migrant (binary, not a migrant/migrant) | Water column verticality: 1 = demersal; 2 = benthopelagic; 3 = pelagic |
| Dragonflies | Hindwing length (mm) | Larval size (mm) | number of eggs | 1 = less than a year; 2 = more than a year | - | 1 = carnivore | Residuals of $\log(\text{hindwing length})$ and $\log(\text{abdomen length})^{18}$ | - |
| Butterflies | Forewing length (mm) | Egg size (mm) | Number of eggs | 1 = one generation per year; 2 = two generations per year | 1 = monophagous; 2 = oligophagous; 3 = polyphagous | 1 = herbivore | - | - |
| Grasshoppers | Total length (mm) | - | Number of eggs | 1 = less than one year; 2 = more than a year | - | - | 1 = flightless, 2 = dimorphic, 3 = winged | 1 = troglobiont; 2 = terricolous; 3 = graminicolous/forbicolous; 4 = arbusticolous; 5 = arboricolous |
| Spiders | Body length (mm) | - | - | Generation length (years) | 1 = stenophagous; 2 = euriphagous | 1 = carnivore | Ballooning frequency: 1 = rare; 2 = occasional; 3 = frequent | Verticality index (Macias-Hernandez et al. 2020) |
| Snails | Heometric mean of the length and width of a shell (mm)<sup>49</sup> | Egg diameter (mm) | 1 = 1-10 eggs; 2 = 10-100 eggs | 1 = 1-2 years; 2 = 2-5 years; 3 = >5 years | Number of classes eaten. Classes = detritus/litter/living material/dead material/herbivore/carnivore | 1 = non-carnivore; 2 = carnivore | - | 0 = caves and other subterranean habitats; 1 = under-rocks, 2 = rock level; 3 = above rocks; 4 = vegetation |
| Bryophytes | Shoot length (mm) | Spore diameter (um) | - | 1 = annual or biannual, 2 = perennial | - | 1 = produce | 1 / $\log(\text{spore size})^{21}$ | 1 = exclusively soil/rock; 2 = trees/walls/tree logs |
| Ferns | Height<br>(cm) | - | - | - | - | 1 =<br>produce<br>r | - | 1 = terrestrial/aquatic; 2<br>= soil or epiphytes; 3 =<br>epiphytes/trunk |
| Gymnosperms | Height<br>(cm) | Seed max<br>diameter<br>(mm) | - | Generation<br>length<br>(years) | - | 1 =<br>produce<br>r | - | - |
| Monocots | Height<br>(cm) | Seed<br>weight<br>(Thousand kernel<br>weight)<br>(g) | - | Generation<br>length<br>(years) | - | 1 =<br>produce<br>r | - | - |
| Legumes | Height<br>(cm) | Seed max<br>diameter<br>(mm) | - | Generation<br>length<br>(years) | - | 1 =<br>produce<br>r | - | - |

**Table S4:** Proxies for Habitat breadth, Altitudinal range, Human footprint and Change in human footprint. All data extracted or derived from the IUCN Red List database.

| Habitat breadth | Altitudinal range | Human footprint<br>(HFP) | Change in human<br>footprint | Geographical<br>range size |
| --- | --- | --- | --- | --- |
| Number of<br>habitat types | Maximum -<br>minimum<br>elevation | Mean human<br>footprint across<br>geographical range | Mean change in<br>HFP values across<br>geographical range | Extent of<br>Occurrence<br>(km<sup>2</sup>). |

**Table S5:** Number of species per taxonomic group and biogeographic realm.

|  | Afrotropic | IndoMalay | Nearctic | Neotropic | Australasia | Paelearctic |
| --- | --- | --- | --- | --- | --- | --- |
| **Mammals** | 10 | 10 | 10 | 10 | 10 | 10 |
| **Birds** | 10 | 10 | 10 | 10 | 10 | 10 |
| **Reptiles** | 10 | 10 | 10 | 10 | 10 | 10 |
| **Amphibians** | 10 | 10 | 10 | 10 | 10 | 10 |
| **Fishes** | 10 | 10 | 10 | 10 | 10 | 10 |
| **Dragonflies** | 10 | 10 | 10 | 10 | 10 | 10 |
| **Butterflies** | 10 | 2 | 0 | 16 | 16 | 16 |
| **Grasshoppers** | 30 | 0 | 0 | 0 | 0 | 30 |
| **Spiders** | 0 | 0 | 0 | 0 | 0 | 39 |
| **Snails** | 0 | 0 | 0 | 0 | 0 | 60 |
| **Bryophytes** | 0 | 0 | 0 | 0 | 0 | 60 |
| **Ferns** | 10 | 10 | 10 | 10 | 10 | 10 |
| **Gymnosperm**<br>**s** | 10 | 10 | 10 | 10 | 10 | 10 |
| **Monocots** | 10 | 10 | 10 | 10 | 10 | 10 |
| **Fabales** | 10 | 10 | 10 | 10 | 10 | 10 |

## Supplementary Figures

**Figure S1:**
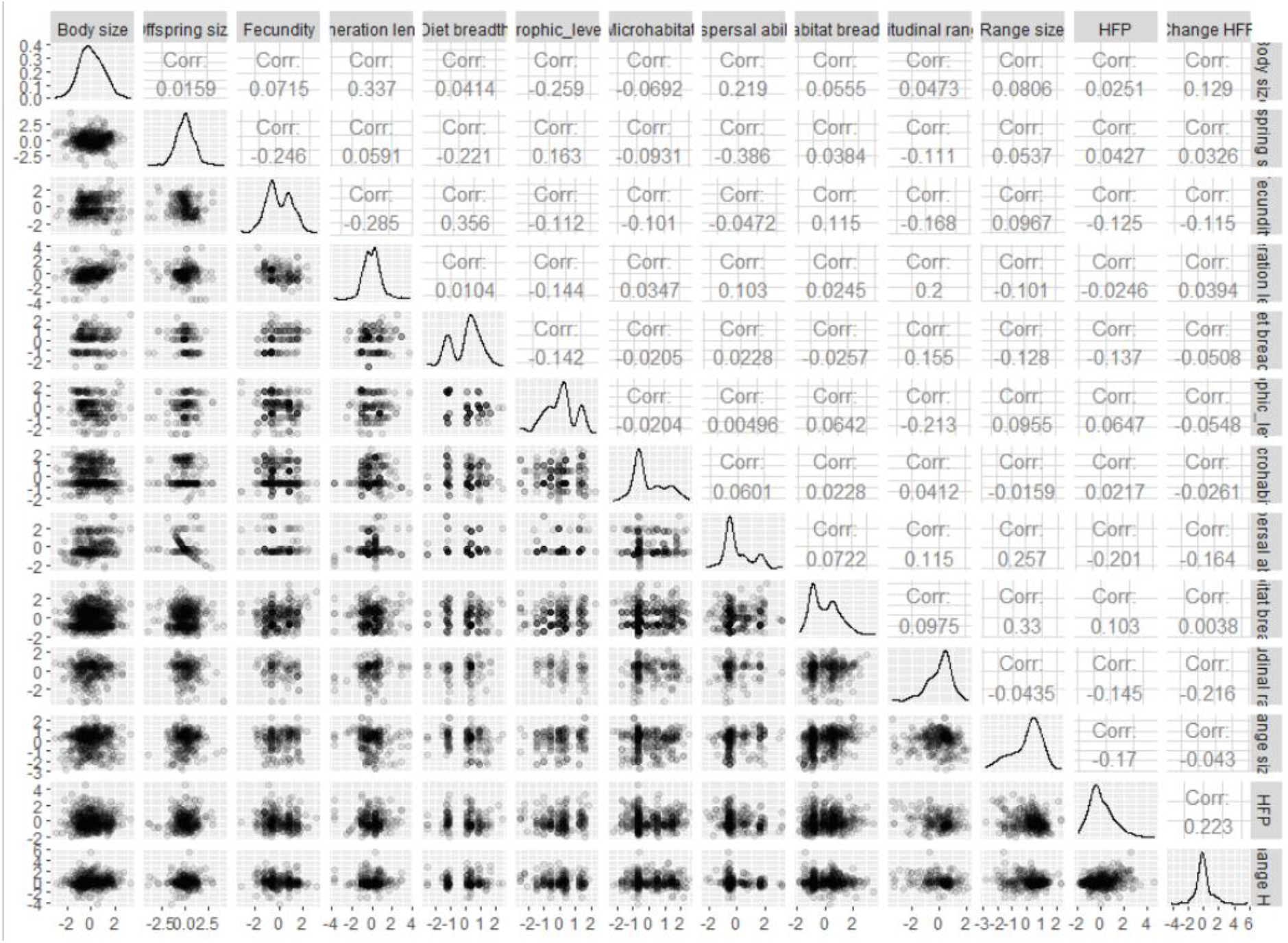
Pairwise correlations between traits. Upper panel: Spearman’s rank correlation coefficient between each pair of traits. Diagonal: histogram of each trait. Lower panel: scatterplots of each trait. Higher density of datapoints is indicated by darker shades of gray.

**Figure S2:**
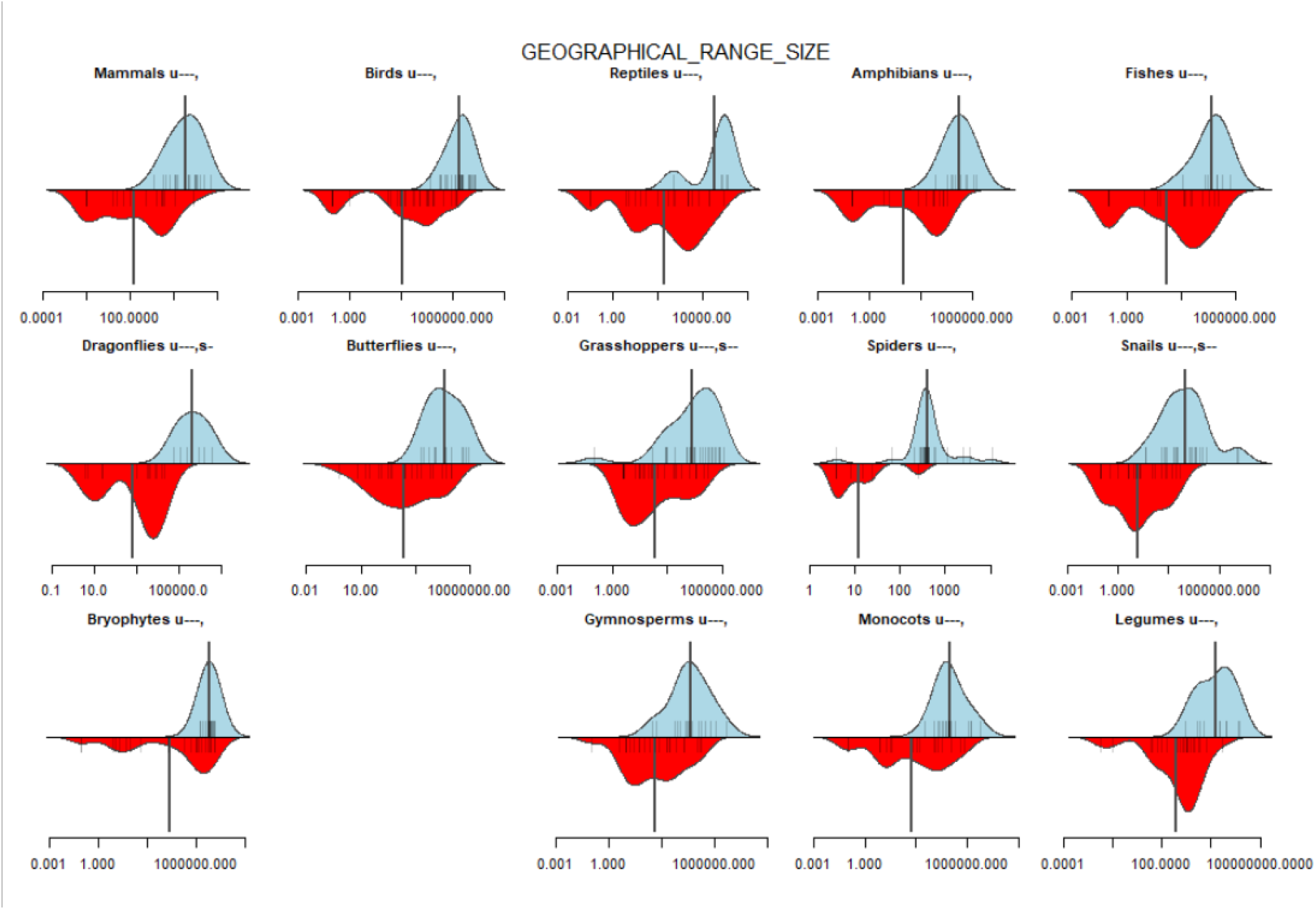
Beanplots of density of geographical range size values between non-threatened (blue, upper side) and threatened (red, lower side) species. Small vertical bars represent one species’ value; darker bars indicate several species with the same trait value. The large vertical bar is the mean geographical range size value. Null models show whether the mean (u) or standard deviation (s) of threatened species is higher (+++, ++, +) or lower (−−−, −−, −) than expected. Significance codes: +++ or −−− x < 0.01; ++ or −− 0.01 <= x < 0.05; + or - 0.05 <= x < 0.1. The Bayesian model was highly significant (pMCMC < 0.0001, N = 506).

## Author contributions

FC and PC conceptualized the initial idea of the paper. All authors contributed to the acquisition of the data. FC conducted the formal analysis. FC and PC wrote the first draft of the manuscript, and all authors read, and contributed with suggestions.

## Acknowledgments

F.C. and P.C. were funded by Kone Foundation, Finland, with the project ‘Trait-based prediction of extinction risk’.

## References

1. Vos, J. M. D., Joppa, L. N., Gittleman, J. L., Stephens, P. R. & Pimm, S. L. Estimating the normal background rate of species extinction. Conserv. Biol. 29, 452–462 (2015).

2. Purvis, A., Gittleman, J. L., Cowlishaw, G. & Mace, G. M. Predicting extinction risk in declining species. Proc. R. Soc. Lond. B Biol. Sci. 267, 1947–1952 (2000).

3. Marco, M. D., Venter, O., Possingham, H. P. & Watson, J. E. M. Changes in human footprint drive changes in species extinction risk. Nat. Commun. 9, 1–9 (2018).

4. Chichorro, F., Juslén, A. & Cardoso, P. A review of the relation between species traits and extinction risk. Biol. Conserv. 237, 220–229 (2019).

5. Cardillo, M. et al. Human Population Density and Extinction Risk in the World’s Carnivores. PLoS Biol 2, e197 (2004).

6. Cardoso, P., Erwin, T. L., Borges, P. A. V. & New, T. R. The seven impediments in invertebrate conservation and how to overcome them. Biol. Conserv. 144, 2647–2655 (2011).

7. Verde Arregoitia, L. D. Biases, gaps, and opportunities in mammalian extinction risk research. Mammal Rev. 46, 17–29 (2016).

8. Bennett, P. M. & Owens, I. P. Variation in extinction risk among birds: chance or evolutionary predisposition? Proc. R. Soc. Lond. B Biol. Sci. 264, 401–408 (1997).

9. Gage, G. S., Brooke, M. de L., Symonds, M. R. E. & Wege, D. Ecological correlates of the threat of extinction in Neotropical bird species. Anim. Conserv. 7, 161–168 (2004).

10. Nolte, D., Boutaud, E., Kotze, D. J., Schuldt, A. & Assmann, T. Habitat specialization, distribution range size and body size drive extinction risk in carabid beetles. Biodivers. Conserv. 28, 1267–1283 (2019).

11. Venter, O. et al. Global terrestrial Human Footprint maps for 1993 and 2009. Sci. Data 3, 160067 (2016).

12. Böhm, M. et al. Correlates of extinction risk in squamate reptiles: the relative importance of biology, geography, threat and range size. Glob. Ecol. Biogeogr. 25, 391–405 (2016).

13. Slatyer, R. A., Hirst, M. & Sexton, J. P. Niche breadth predicts geographical range size: a general ecological pattern. Ecol. Lett. 16, 1104–1114 (2013).

14. Rabinowitz, D. Seven forms of Rarity. in Biological aspects of rare plant conservation 205–217 (John Wiley & Sons Ltd., 1981).

15. Reinhardt, K., Köhler, G., Maas, S. & Detzel, P. Low dispersal ability and habitat specificity promote extinctions in rare but not in widespread species: the Orthoptera of Germany. Ecography 28, 593–602 (2005).

16. Benscoter, A. M. et al. Threatened and Endangered Subspecies with Vulnerable Ecological Traits Also Have High Susceptibility to Sea Level Rise and Habitat Fragmentation. PLoS ONE 8, e70647 (2013).

17. Mattila, N., Kaitala, V., Komonen, A., PäIvinen, J. & Kotiaho, J. S. Ecological correlates of distribution change and range shift in butterflies: Distribution decline in butterflies. Insect Conserv. Divers. 4, 239–246 (2011).

18. MacLean, S. A. & Beissinger, S. R. Species’ traits as predictors of range shifts under contemporary climate change: A review and meta-analysis. Glob. Change Biol. 23, 4094–4105 (2017).

19. Pimm, S. L., Jones, H. L. & Diamond, J. On the Risk of Extinction. Am. Nat. 132, 757–785 (1988).

20. González-Suárez, M., Gómez, A. & Revilla, E. Which intrinsic traits predict vulnerability to extinction depends on the actual threatening processes. Ecosphere 4, 1–16 (2013).

21. Saar, L., Takkis, K., Pärtel, M. & Helm, A. Which plant traits predict species loss in calcareous grasslands with extinction debt? Divers. Distrib. 18, 808–817 (2012).

22. Botts, E. A., Erasmus, B. F. N. & Alexander, G. J. Small range size and narrow niche breadth predict range contractions in South African frogs. Glob. Ecol. Biogeogr. 22, 567–576 (2013).

23. McKinney, M. L. & Lockwood, J. L. Biotic homogenization: a few winners replacing many losers in the next mass extinction. Trends Ecol. Evol. 14, 450–453 (1999).

24. Seppälä, S. et al. Species conservation profiles of a random sample of world spiders IV: Scytodidae to Zoropsidae. Biodivers. Data J. 6, e30842 (2018).

25. Brummitt, N. A. et al. Green Plants in the Red: A Baseline Global Assessment for the IUCN Sampled Red List Index for Plants. PLOS ONE 10, e0135152 (2015).

26. Weiss, K. C. B. & Ray, C. A. Unifying functional trait approaches to understand the assemblage of ecological communities: synthesizing taxonomic divides. Ecography 42, 2012–2020 (2019).

27. Hadfield, J. D. MCMC methods for multi-response generalized linear mixed models: The MCMCglmm R package. J. Stat. Softw. 33, 1–22 (2010).

28. R Core Team. R: A language and environment for statistical computing. (2019).

29. Kampstra, P. Beanplot: A boxplot alternative for visual comparison of distributions. J. Stat. Softw. Code Snippets 28, 1–9 (2008).

30. Wickham, H. ggplot2: Elegant graphics for data analysis. (Springer-Verlag New York, 2016).

31. Schloerke, B. et al. GGally: Extension to ‘ggplot2’. (2020).

32. Olden, J. D., Hogan, Z. S. & Zanden, M. J. V. Small fish, big fish, red fish, blue fish: size-biased extinction risk of the world’s freshwater and marine fishes. Glob. Ecol. Biogeogr. 16, 694–701 (2007).

33. Cardillo, M. et al. Multiple Causes of High Extinction Risk in Large Mammal Species. Science 309, 1239–1241 (2005).

34. Ripple, W. J. et al. Extinction risk is most acute for the world’s largest and smallest vertebrates. Proc. Natl. Acad. Sci. 201702078 (2017) doi:10.1073/pnas.1702078114.

35. Moles, A. T. et al. A Brief History of Seed Size. Science 307, 576–580 (2005).

36. Boyles, J. G. & Storm, J. J. The Perils of Picky Eating: Dietary Breadth Is Related to Extinction Risk in Insectivorous Bats. PLoS ONE 2, e672 (2007).

37. Price, S. A. & Gittleman, J. L. Hunting to extinction: biology and regional economy influence extinction risk and the impact of hunting in artiodactyls. Proc. R. Soc. B Biol. Sci. 274, 1845–1851 (2007).

38. Carvalho, J. C. & Cardoso, P. Drivers of beta diversity in Macaronesian spiders in relation to dispersal ability. J. Biogeogr. 41, 1859–1870 (2014).

